# FuzzyVariantsExplorer: Graded exploration of genomic data and fuzzy queries

**DOI:** 10.1101/811695

**Authors:** Marco Falda

## Abstract

**Summary:** FuzzyVariantExplorer is a web interface and a fuzzy search system for exploring richly annotated genomes in a flexible way. The application allows combining vague constraints in expressive logical queries to retrieve graded sets of genes and their associated variants. Results can be further refined in a visual way by selecting keywords and categories that are extracted from the underlying annotations and weighted with a statistical significance score; keywords can then be clustered using Latent Dirichlet Allocation and explored visually. The system has been applied to a set of *Saccharomyces cerevisiae* genome annotations and a few variants.

**Availability:** FuzzyVariantExplorer source code is available at the URL https://bitbucket.org/mfalda/fuzzyvariantsexplorer/src/ and has been developed in Scala, TypeScript and R.

## I. Introduction

Imprecise knowledge is frequent in life sciences due, for example, to the incomplete understanding of biological mechanisms, the uncertainty about normality ranges for test results, the simultaneous presence of more than one condition or missing information [1].

Traditional database models are not accurate in representation and manipulation of such knowledge. Fuzzy logic has been proposed since [2] for enriching classical data models and allow for processing with uncertain and imprecise data. The same setting has also been proposed for modeling fuzzy databases [3]. Life sciences need such sophisticated models for manipulating complex objects and semantic relationships arising from the vague entities involved in their processes.

The system described in this paper, named FuzzyVariantExplorer, does not try to add another theoretical setting for modeling data in a fuzzy way, but uses a simple and minimal layer for expressing queries in the database in a more flexible way. Namely, it allows exploring annotated genomes stored in traditional relational database management systems (RDBMSs) by expressing a preference degree on various criteria used to formulate the queries. This means that the user can be less accurate when searching terms and expressing numerical information, and that a possibility degree will be associated to the results according to her initial preferences.

Additionally, results are summarized in a graphical way according to different criteria, and the statistical significance of the resulting categories is estimated. A Latent Dirichlet Allocation (LDA)[4] can be performed on each criteria in order to obtain a more general idea of the collected results.

## II. Theory

### A. Fuzzy Logic

According to Zadeh [2], real world objects often do not present a crisp membership; this is why classical logics has difficulties to describe some knowledge (e.g. the concepts of “tall”, “young”, etc.). Another problem is that common information is affected by imprecision or vagueness.

He defined the notion of membership degree which allows to express how much an element belongs to a set; this degree is defined in the range [0, 1]. A fuzzy set *F* is defined on the universe *U* by means of a membership function *µ*_*F*_ that gives the degree of membership *µ*_*F*_ (*u*) ∈ [0, 1] of an element *u* ∈ *F*. Depending on the application a membership degree can be interpreted as:

- a similarity degree with respect to the typical members of the fuzzy set;
- a preference degree that grade the elements of the fuzzy set;
- an uncertainty degree which tells the possibility that an element belongs to the fuzzy set.

The set of the elements which are fully possible, that is *C*_*F*_ (*u*) = *u* ∈ *U* : *µ*_*F*_ (*u*) = 1 is called “core” of *F*; the “support” of *F* is defined as the characteristic function of *F*, that is *χ*_*F*_ (*u*) = *u* ∈ *U* : *µ*_*F*_ (*u*) > 0.

An alternative definition of a fuzzy set *F* can be given in terms of a set of sets associated to a threshold on the original set. These sets are the *α*-cuts of *F* and are defined as

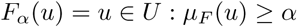

If the universe *U* is equal to the real numbers set, each fuzzy set defined on *U* is called a “fuzzy quantity”. A membership function which is semi-continuous and has only a fully possible element is called a “fuzzy number”.

#### Operations

The classical operations on sets are extended as follows:

- *F* is included in *G* if

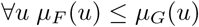
- the complement of *F* in *U*, denoted as 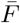 is

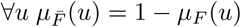
- the intersection between *F* and *G* is

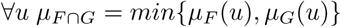
- the union between *F* and *G* is

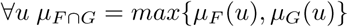

These extensions are not unique, they obey to the so-called “max-min” theory. In this theory two properties of the classical Boolean algebra are not satisfied: the law of the excluded middle and the law of non contradiction. These two exceptions are recovered when *F* and *G* are classical sets.

The exclusive or can be easily derived from its logical equivalence: *F*⊕*G* is equivalent to *F* ∨*G* ∧¬(*F* ∧*G*), therefore:

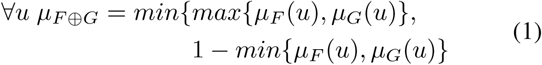

The logical implication has been defined in several ways in the fuzzy logic field, for example:

- Lukasiewicz: ∀*u min*{1, 1 − *µ*_*F*_ (*u*) + *µ*_*G*_(*u*)}
- Kleene-Dienes: ∀*u max*{1 − *µ*_*F*_ (*u*), *µ*_*G*_(*u*)}
- Mamdani: ∀*u min*{*µ*_*F*_ (*u*), *µ*_*G*_(*u*)}

The latter has a more intuitive meaning for rule sets and has been selected.

### B. Fuzzy SQL

Expressions are processed through one-token look-ahead left-to-right rightmost derivation parsers LARL(1) [5] and transformed in traditional SQL queries enriched with fuzzy operators (see Subsection III-B about User Defined Functions or UDFs). As depicted in Figure 2a, the query is optimized by first filtering on the support of the formula and then applying the fuzzy operators. Three parsers have been defined that correspond to the three optimization steps above: the logical parser for the support, the logical threshold parser for *α*-cuts and the fuzzy parser for translating fuzzy formulas in SQL expressions and UDFs.

**Fig. 1.**
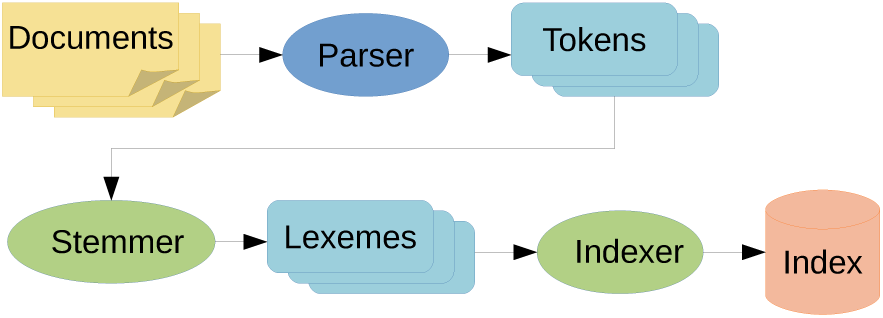
Steps for full-text search indexing in PostgreSQL.

**Fig. 2.**
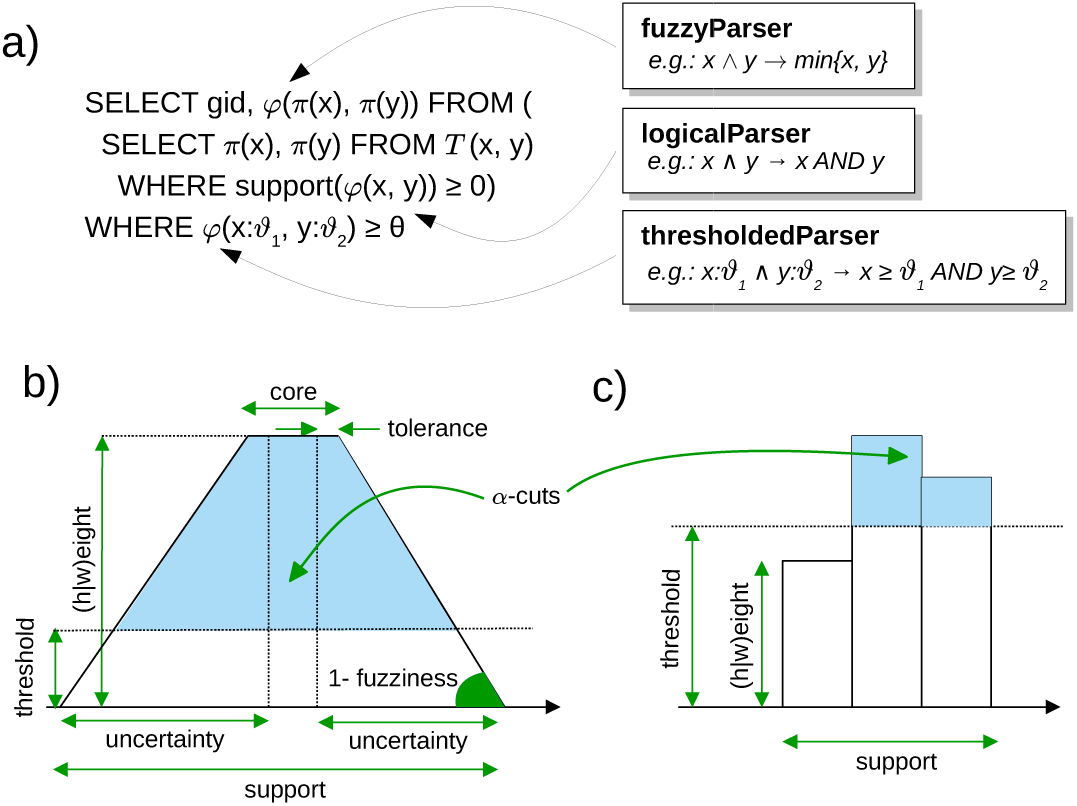
a) the LARL(1) parsers involved in the mapping from a fuzzy query to SQL. b) a trapezoidal t-norm for numerical data. b) a possibility distribution for categorical data.

There are currently three main classes of operators, one for each class of data: fuzzy order operators for numerical data, fuzzy choice operators (discrete possibility distributions) for categorical data and operators for mapping scores into fuzzy preference degrees, typically used for textual data. Numerical operators are based on trapezoidal t-norms [6], for example a fuzzy interval is given by the following function (Figure 2b):

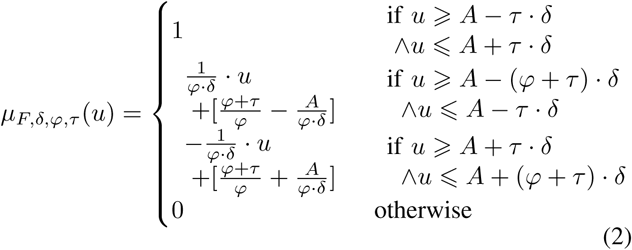

where *δ* represents the uncertainty, *φ* the fuzziness of the statement and *τ* the tolerance.

Categorical data are modeled as a graded set, that is a set in which each element is associated with a preference degree (Figure 2c). Textual data are retrieved using the sophisticated PostgreSQL full text search (FTS) facility [7] (Figure 1): documents are segmented in tokens and then these are reduced to a form common to all word variations (stemming) in order to increase search effectiveness; finally, the resulting lexemes are indexed in the database using a generalized inverted index.

To ensure the equivalence between fuzzy and classical operators whenever the *α*-cuts set is reduced to a binary set, the following function is applied to the scores coming from categorical and textual data:

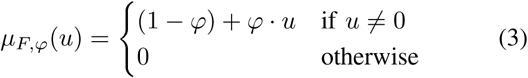

In the case of the two latter data types, the tolerance is more difficult to model because it can be seen as a sort of imprecision on the uncertainty intervals. It could be treated as suggested in [8], where an orthogonal idea of “coarseness” is combined with the fuzzy preference degrees: tolerance could be seen as an uncommitted choice among neighboring concepts or categories, for example “canine” would mean “dog” or “wolf” or “fox” *et c.* with a lower preference degree depending on an information content metric.

### C. Natural Language Processing

Simply taking single words could miss some interesting characterizations given by the association of adjectives or attributive nouns, the so called “nominal phrases” (NP); for this reason all words in the textual data are first categorized according to their part of speech (POS tagging) using the Stanford NLP toolkit [9] in Scala.

Tags belong to the Penn Treebank English POS tag set [10] and, as a first simple criterion, types “NN”, “NNP”, “NNS” and “JJ” have been aggregated together.

## III. Implementation

FuzzyVariantExplorer is a web application, developed in Scala using the Play2 MVC framework. TypeScript has been adopted to increase the safeness and the scalability of client side code; the application is organized in three intuitive sections: the query interface, the results page and the reports for individual genes or variants. The results page presents three sub-sections containing a table with the matching genes, a set of charts and word clouds for a visual overview and a graphical interface for the LDA topics (see Subsection III-D).

In the first section queries can be composed by the user in a visual way by choosing from a hierarchical menu of criteria. Criteria are incrementally added to the formula by means of logical connectives and can be negated or moved into inner priority levels. More than one query can be built and it will be executed concurrently in order to allow for successive comparisons. Another advantage of using two levels of formulas is to pre-compute possibly over-constrained criteria, that would otherwise get no results, and in a later step inspect the conflicting sets.

### A. A schema for INI files

FuzzyVariantExplorer interface has been designed to be quite configurable as far as new criteria and statistics are concerned. To add new criteria it is in fact sufficient to provide database views that link the data with the IDs of genes (or whatever entity the application is currently applied at) and the fields with the criteria. The same goes for new report sections and statistics.

Once these new views have been provided, they have to be notified to the system by writing new sections in a INI file, which is a very simple and clear file format. However, a limit of the INI file format is that there is not a way to check its consistency by means of a schema, like for example in the case of XML. For this reason, a INI validator has been created in Scala and then integrated in FuzzyVariantExplorer; it is able to identify spurious or missing keys and also to ensure the existence of schemata, tables and fields in the database. A general standalone version has been implemented in Rust language [11].

### B. User Defined Functions in PL/pgSQL

In order to express in SQL the possibility distributions, seven user defined functions for PostgreSQL have been written (see Table I). They have been defined using the integrated PL/pgSQL language, since their implementation in C as shared objects did not offer a significant increase in performance, therefore a more flexible way of development has been preferred.

**TABLE I.** User Defined Functions added to PostgreSQL.

| Name | Distribution |
| --- | --- |
| fuzzy_pos(x, center, delta, fuzziness, tol) | trapezoidal |
| fuzzy_lt(x, center, delta, fuzziness, tol) | < |
| fuzzy_le(x, center, delta, fuzziness, tol) | ≤ |
| fuzzy_ne(x, center, delta, fuzziness, tol) | ≠ |
| fuzzy_ge(x, center, delta, fuzziness, tol) | ≥ |
| fuzzy_gt(x, center, delta, fuzziness, tol) | > |
| fuzzy_distr(cat, preferences, fuzziness) | discrete |

### C. Compile-time query generation with Quill

Several SQL queries in the source code have been written in Quill [12] a quoted domain specific language created by a Twitter software engineer. It has been inspired by a paper about language-integrated queries [13] and it allows for compile-time query generation and asynchronous execution, a modality well accepted in the Play2’s share-nothing architecture based on actors.

The model definition is simply a Scala case class:

~~~
**case class User**(id**: Long**, name**: String**,
    isActive**: Boolean**)
~~~

Queries are prepared by quoting Scala-like expressions

~~~
**def** byId(id**: Long**) **=** quote {
    **Users**.filter(**_**.id == lift(id))
}
~~~

and execution returns a future:

~~~
**def** find(id**: Long**)**: Future**[**Option**[**User**]] **=**
    db.run(byId(id)).map(**_**.headOption)
~~~

### D. Statistical tests and Rserve

The implemented tool has been designed to be used with huge genomic data, therefore a useful feature is to filter the most meaningful results by applying statistical tests to them. The general principle underlying all types of data (categorical, textual, numerical) is to compute the frequency of all labels, nominal phrases or intervals in the data associated to a given criterion and then determine the p-values for the filtered entities using a proportions test adjusted by a Benjamini-Hochberg method [14].

To have an even more abstract view of the results, LDA [4] is applied and the topics are then rendered using the interactive LDAvis tool, which provides a global view and how they differ from each other, while at the same time allowing for a deep inspection of the terms most highly associated with each individual topic [15].

Since all these statistical tests and procedures have already been provided in R packages, they have been implemented as R functions and called by sending them to Rserve [16].

## IV. Example application

The system has been applied to a set of *Saccharomyces cerevisiae* genome annotations collected from the SGD Project [17] using programs written in GO language [18]. Several types of data have been collected about the cited sequences: numerical data regarding the starting and ending coordinates in the chromosomes, molecular weights, isoelectric points; categorical data about protein domains and ontological terms; textual data referring to several descriptions.

By searching the word “tubulin” in gene descriptions, the word clouds shown in Figure 4, Figure 5 and Figure 6 have been obtained from descriptions of Biological Process (BP), Molecular Function (MF) and Cellular Component (CC) sub-ontologies respectively. In the left parts of each figure there is the unfiltered distribution, in which it is easy to recognize the most typical terms of each sub-ontology. On the other hand, in the right parts of the figures there are more specific terms that the system has been able to extract taking into account the original query, such as “microtubules” in BP, “actin” in MF and “tubulin” in CC.

**Fig. 3.**
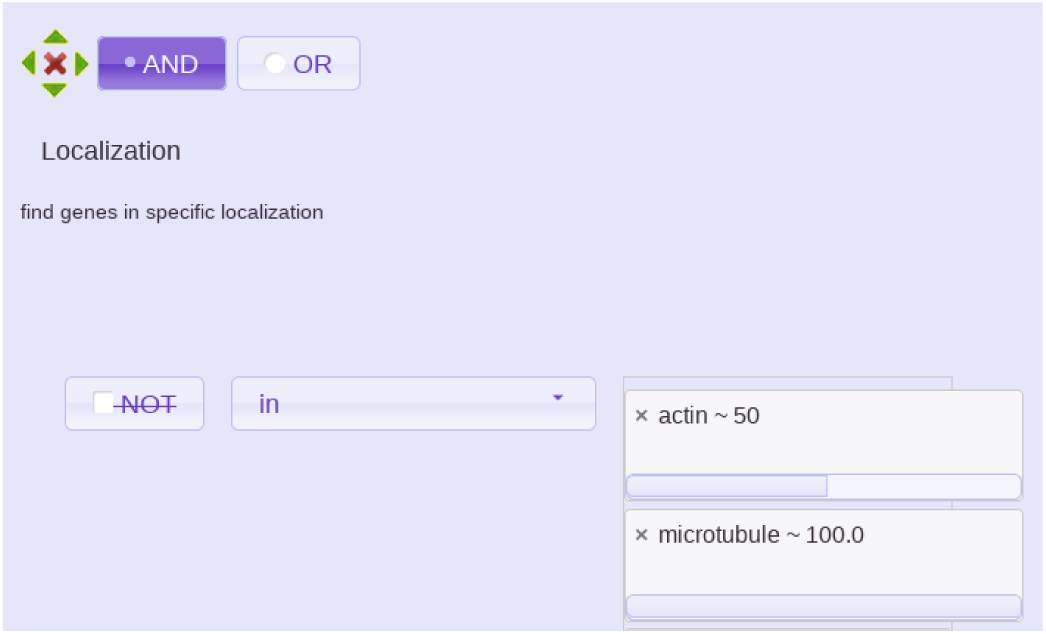
Example of a criteria for categorical data: “microtubule” is preferred w.r.t. “actin”.

**Fig. 4.**
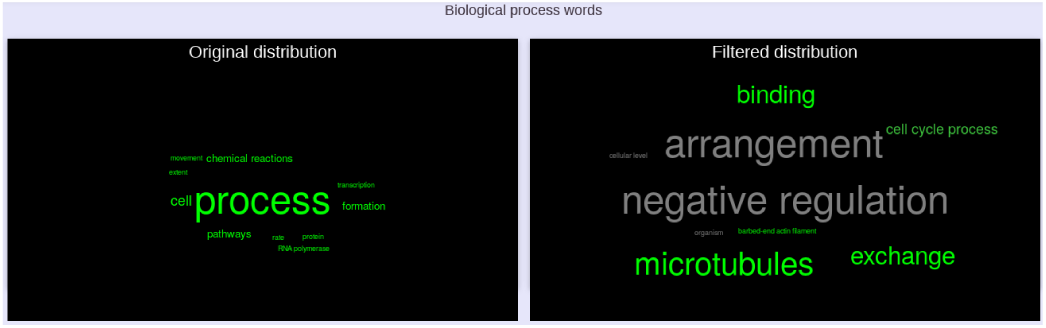
Word cloud for Biological Process sub-ontology.

**Fig. 5.**
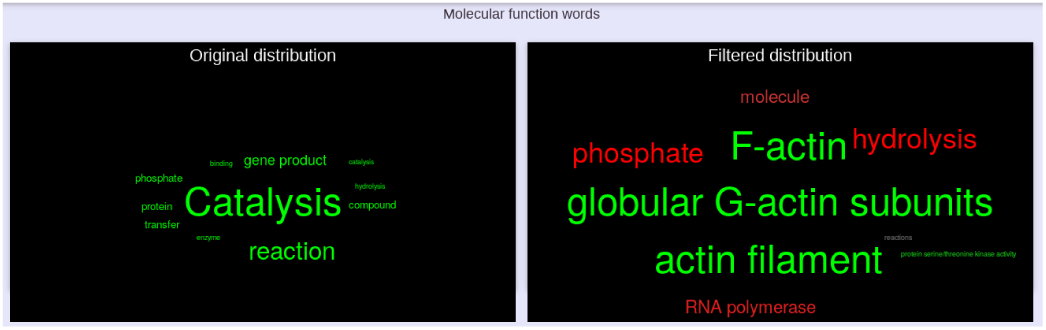
Word cloud for Molecular Function sub-ontology.

**Fig. 6.**
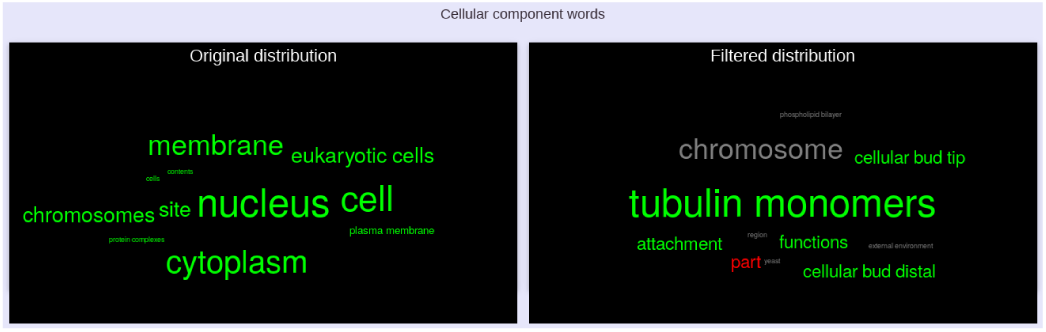
Word cloud for Cellular Component sub-ontology.

In Figure 7 the LDA analysis on words belonging to gene descriptions has been reported; this analysis tries to cluster the entities in meaningful concepts, for example cluster n.3 is labeled by words like “Alphatubulin”, “F-actin” and so on.

**Fig. 7.**
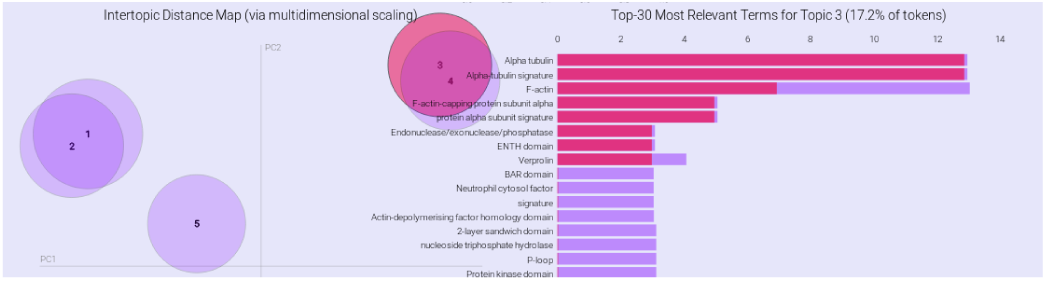
Latent Dirichlet Analysis of gene descriptions.

## V. Conclusion

Modern genomic annotation systems rely on huge amounts of data, therefore to ensure a rapid access and a maintainable infrastructure the traditional RDBMSs are still a valid solution.

Independently of the underlying data storage model, database systems are designed with the assumption that precise information will be stored. Whenever the knowledge of the reality to be modeled is imperfect their power becomes less relevant. In such cases tools for describing uncertain or imprecise information have to be applied.

FuzzyVariantExplorer provides a novel query interface that on one hand allows for composing complex queries in a simple way and on the other hand adds more flexibility by exploiting fuzzy operators. Moreover, since genomic annotation data are vast, results can be summarized graphically and the most relevant topics extracted for several criteria.

## Notes

https://bitbucket.org/mfalda/fuzzyvariantsexplorer/src/default/

